# Premature endocycling of *Drosophila* follicle cells causes pleiotropic defects in oogenesis

**DOI:** 10.1101/2023.10.10.561736

**Authors:** Hunter C. Herriage, Brian R. Calvi

**Affiliations:** Department of Biology, Indiana University, Bloomington, IN 47405; Melvin and Bren Simon Cancer Center, Indianapolis, IN; Indiana University School of Medicine, Bloomington, IN

**Keywords:** endocycling, border cells, polyploidy, oogenesis, epithelium, *Drosophila*

## Abstract

Endocycling cells grow and repeatedly duplicate their genome without dividing. Cells switch from mitotic cycles to endocycles in response to developmental signals during the growth of specific tissues in a wide range of organisms. The purpose of switching to endocycles, however, remains unclear in many tissues. Additionally, cells can switch to endocycles in response to conditional signals, which can have beneficial or pathological effects on tissues. However, the impact of these unscheduled endocycles on development is underexplored. Here, we use *Drosophila* ovarian somatic follicle cells as a model to examine the impact of unscheduled endocycles on tissue growth and function. Follicle cells normally switch to endocycles at mid-oogenesis. Inducing follicle cells to prematurely switch to endocycles resulted in lethality of the resulting embryos. Analysis of ovaries with premature follicle cell endocycles revealed aberrant follicular epithelial structure and pleiotropic defects in oocyte growth, developmental gene amplification, and the migration of a special set of follicle cells known as border cells. Overall, these findings reveal how unscheduled endocycles can disrupt tissue growth and function to cause aberrant development.

**Summary Statement:** A premature switch to polyploid endocycles in *Drosophila* ovarian follicle cells caused pleiotropic defects in oogenesis and compromised female fertility, revealing new ways in which unscheduled endocycles cause developmental defects.

## Introduction

Many tissues grow via a variant cell cycle known as the endocycle. Endocycles are characterized by alternating G and S phases without mitotic division, resulting in the growth of large polyploid cells (Calvi 2013; Ovrebo and Edgar 2018; Fox *et al*. 2020). A regulated switch from mitotic cycles to endocycles is part of the normal developmental growth programs of specific tissues across *Eukarya* (Ovrebo and Edgar 2018; Fox *et al*. 2020). In contrast, cells can also switch to unscheduled endocycles in response to stress and other conditional signals (Hassel *et al*. 2014; Rotelli *et al*. 2019; Bailey *et al*. 2021; Besen-McNally *et al*. 2021). These induced endocycling cells can be beneficial for wound healing and regeneration, but they can also have pathological effects, including the promotion of genome instability and cancer (Shu *et al*. 2018; Stormo and Fox 2019; Bailey *et al*. 2021; Almeida Machado Costa *et al*. 2022; Pienta *et al*. 2022; Herriage *et al*. 2023). Much remains unknown, however, about how unscheduled endocycling affects tissue structure and function.

In this study we used the somatic follicle cells of the *Drosophila* ovary as a model to evaluate the effects of unscheduled endocycles on tissue function. A *Drosophila* ovary consists of ∼16 ovarioles, each composed of one germarium stem cell compartment followed by an “assembly line” of progressively developing egg chambers in 14 stages of oogenesis (Fig. 1A) (King 1970; Duhart *et al*. 2017). These developing egg chambers contain the germline (15 endocycling nurse cells and one oocyte) ensheathed by an epithelium of somatic follicle cells (Fig. 1A) (King 1970; Jia *et al*. 2015; Duhart *et al*. 2017). Follicle cells originate from a population of two to four somatic stem cells in the germarium and proliferate mitotically over the course of three days until stage 6 of oogenesis (Margolis and Spradling 1995; Sun and Deng 2007; Fadiga and Nystul 2019). In response to Notch signaling from the germline at stages 6 / 7, follicle cells switch to the endocycle and subsequently complete three genome duplications over the next ∼24 hours (Fig. 1A) (Spradling 1993; Calvi *et al*. 1998; Deng *et al*. 2001; Lopez-Schier and St Johnston 2001). After achieving a DNA copy number (C) of 16C, follicle cells asynchronously arrest endocycles at different times during late stage 9 (Calvi *et al*. 1998). Follicle cells then synchronously initiate a specialized DNA rereplication program in stage 10B that selectively amplifies the DNA copy number of eggshell (chorion) protein genes, a process known as chorion gene amplification (Fig. 1A) (Spradling and Mahowald 1980; Calvi *et al*. 1998). Disruption of this developmental gene amplification causes thin eggshells and embryonic lethality (Spradling and Mahowald 1980; Calvi 2006; Hua and Orr-Weaver 2017). It has been shown that the endocycle to amplification transition requires the downregulation of Notch signaling and the modification of several transcription factor and cell cycle protein activities (Calvi *et al*. 1998; Royzman *et al*. 1999; Bosco *et al*. 2001; Cayirlioglu *et al*. 2001; Beall *et al*. 2002; Beall *et al*. 2004; Calvi 2006; Sun *et al*. 2008; Huang *et al*. 2013; Jia *et al*. 2015). Much remains unknown, however, about the molecular mechanism of endocycle arrest and the selective activation of amplification origins at precise times of oogenesis. Follicle cells also support egg growth by mediating transport of yolk proteins into the oocyte and participate in the axial patterning of the egg and embryo (Brennan *et al*. 1982; Raikhel and Dhadialla 1992; Duhart *et al*. 2017; Merkle *et al*. 2020). While it is known that follicle cells have these important roles in oogenesis, it remains unclear what biological function is served by the endocycle switch occurring specifically at stage 6. In this study, we find that inducing endocycles to switch to endocycles earlier than stage 6 has severe negative consequences for oogenesis and female fertility, with broader impacts on understanding how unscheduled endocycles may contribute to human disease.

**Figure 1:**
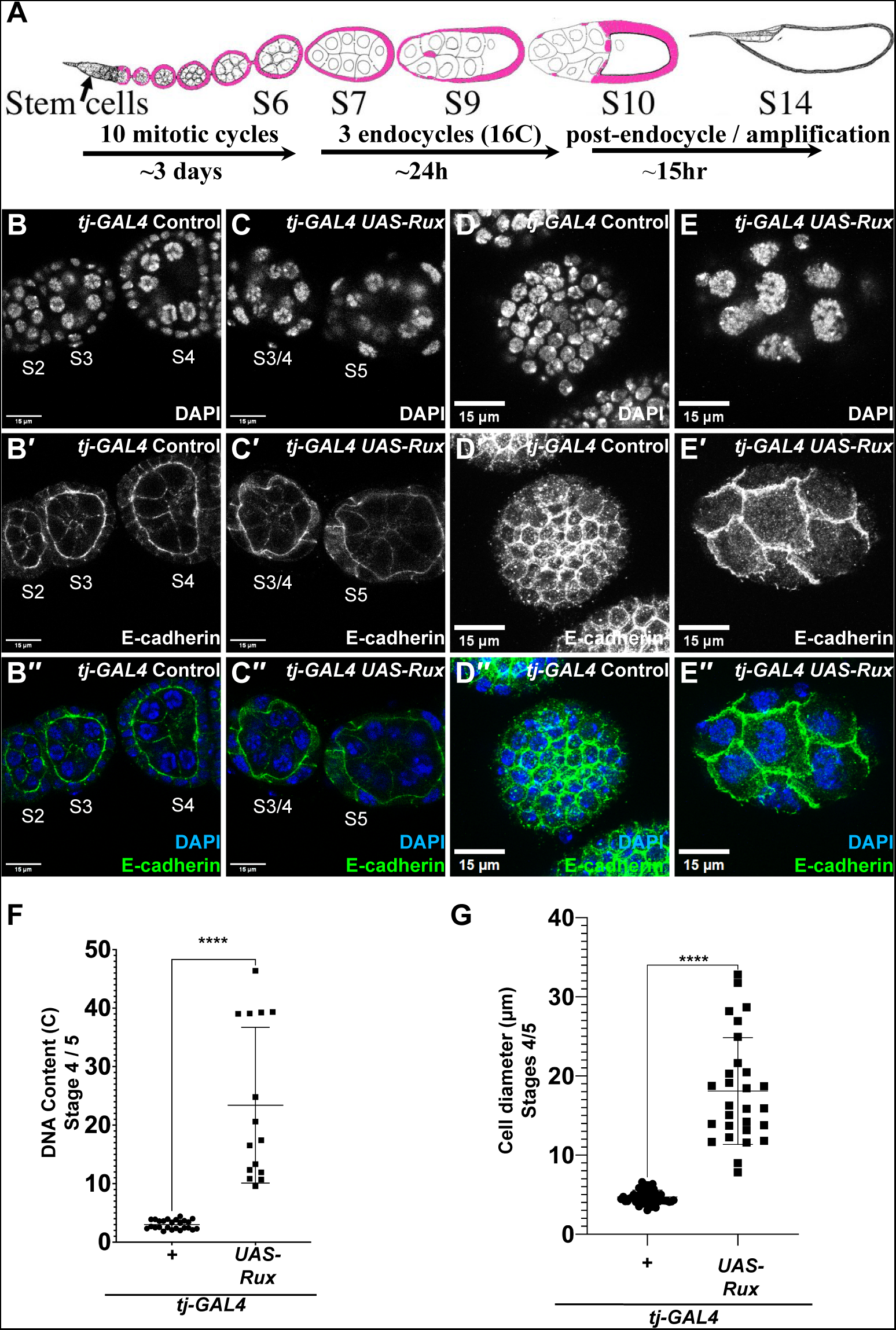
Premature endocycling disrupts early-stage follicular epithelial structure. (A) Drawing of an ovariole with select stages of oogenesis and follicle cell cycle transitions indicated below. Follicle cells are pink. (B-C′′) DAPI staining and immunolabeling against E-cadherin in control (B-B′′) or Rux-expressing (C-C′′) early-stage egg chambers after two days of incubation at 29°C. Stages (S) of oogenesis and germarium (G) are indicated. Scale bar = 15 μm. (D-E′′) DAPI staining and immunolabeling against E-cadherin in cross-section images through the follicle cell epithelium on the surface of control (D-D′′) or Rux-expressing (E-E′′) stage 4 / 5 egg chambers after two days of incubation at 29°C. (F) Quantification of DNA Content (C) from control and Rux-expressing stage 4 / 5 egg chambers after two days of incubation at 29°C; ****p < 0.0001; n = 25 nuclei for control and n = 15 nuclei for *UAS-Rux* across three biological replicates. (G) Quantification of cell diameter from control and Rux-expressing stage 4 / 5 egg chambers after two days of incubation at 29°C. ****p < 0.0001; n = 49 cells for control and n = 28 cells for *UAS-Rux* across three biological replicates.

## Results

### Premature endocycles cause structural defects in the follicular epithelium

To address whether a switch to follicle cell endocycles before stage 6 would affect oogenesis, we induced premature endocycling by expressing the protein Roughex (Rux), a repressor of Cyclin A / CDK1 activity and mitotic entry that has been shown to induce endocycles in other tissues (Thomas *et al*. 1994; Sprenger *et al*. 1997; Avedisov *et al*. 2000; Vidwans *et al*. 2002). We used a *tj-GAL4* transgene to induce expression of a *UAS-Rux* transgene in all ovarian somatic cells, and a temperature-sensitive GAL80^ts^ repressor of GAL4 to control the timing of expression (Brand and Perrimon 1993; McGuire *et al*. 2004; Weaver *et al*. 2020). The *tj-GAL4 tub-GAL80ts / UAS-Rux* experimental and *tj-GAL4 tub-GAL80^ts^ / +* control animals were raised at 18°C to repress GAL4 activity during development. Adult flies were then shifted to 29°C to induce *UAS-Rux* expression and ovaries were examined one and two days thereafter. Given that expression of *Cyclin A* is repressed in endocycling follicle cells after stage 6, it was expected Rux would only impact follicle cell mitotic cycles (Sun and Deng 2005). Consistent with this, within one day after *UAS-Rux* expression, stage 1-6 follicle cells ceased labeling for the mitotic marker phosphorylated Histone H3 (pH3) but continued to incorporate the nucleotide analog 5-ethynyl 2′-deoxyuridine (EdU), indicating that they had switched to G / S endocycles (Fig. S1A-F).

To determine the effects of induced endocycling on early stages of oogenesis, we examined the structure of the follicular epithelium. To do this, we immunolabeled the cell periphery with antibodies against the membrane-associated cell-cell junction protein E-cadherin and stained nuclei with DAPI (Tepass *et al*. 1996; Uemura *et al*. 1996; Oda *et al*. 1997; Godt and Tepass 1998). After two days at 29°C, follicle cells in control ovaries had the normal uniform, cuboidal structure and E-cadherin labeling that was greatest on their apical surface adjacent to the germline (Fig. 1B-B′′) (Godt and Tepass 1998; Gonzalez-Reyes and St Johnston 1998). In contrast, after two days of Rux expression, follicle cells were all larger with wide variations in their size and shape (Fig. 1C-C′′). Some follicle cells had highly irregular shapes, with different parts of the cell having unequal thicknesses in the apical-basal axis, and many with highly irregular E-cadherin labeling (Fig. 1C-C′′). In confocal sections through the middle of egg chambers, the interface between the follicular epithelium and germline appeared wavy instead of the normal smooth circular interface, indicating global deformations of tissue morphology (Fig. 1B′, C′).

Measurement of the DNA content and diameter of the follicle cells on the surface of stage 4 / 5 egg chambers revealed that premature endocycling follicle cells were polyploid and larger than mitotic cycling control follicle cells consistent with their growth through hypertrophy (Fig. 1D-G). These measurements also showed that these premature endocycling cells have a large variance in their size and shape (Fig. 1D-G). Immunolabeling for the cell junction protein beta-catenin, which is known as Armadillo (Arm) in *Drosophila,* or the membrane-associated protein Discs large 1 (Dlg1), also supported the conclusion that premature endocycles caused aberrant cell and epithelial structure and irregular cell-cell junctions (Figs. S2 and S3). Although cell morphology was highly aberrant, Dlg1 had an overall normal basolateral localization, suggesting that there was not a severe disruption of cell polarity (Fig. S3) (Woods *et al*. 1996; Khoury and Bilder 2020). Induction of premature endocycles by RNAi knockdown of *cyclin A* or over-expression of the Cdh1 ubiquitin ligase, *fizzy-related* (*fzr*) also resulted in aberrant cell and epithelial structure, indicating that these effects are not specific to *Rux* overexpression (Fig S4). Overall, these data indicate that induction of unscheduled follicle cell endocycles earlier in oogenesis results in extremely aberrant cell and epithelial structure.

### Premature endocycling increases follicle cell terminal ploidy and causes cell cycle defects in later stages of oogenesis

We next asked what effect premature endocycles would have on follicle cells and egg chambers as they matured to later stages of oogenesis (Fig. 2A). Two days after a shift to 29°C, control stage 10B egg chambers had the normal ∼641 follicle cells and DNA content of ∼16C (Fig. 2B-B′, D, E) (Calvi *et al*. 1998; Sun and Deng 2007). In contrast, the induction of premature endocycles two days earlier in *UAS-Rux* expressing ovaries resulted in stage 10B egg chambers that had four-fold fewer follicle cells (∼179) with a nearly a four-fold increase in average DNA content (mean ploidy ∼56 C) (Fig. 2C-E). These stage 10B follicle cells were larger than normal, consistent with the known scaling of cell size with DNA ploidy (Fig. 2B′, C′) (Tepass *et al*. 1996; Uemura *et al*. 1996; Cadart and Heald 2022). The four-fold decrease in cell number together with a four-fold increase in DNA content suggested that Rux-expressing follicle cells had skipped two mitotic divisions and instead underwent two extra endocycle genome doublings. This conclusion is consistent with the known rate of oogenesis at 29°C (Gandara and Drummond-Barbosa 2022). These stage 10B egg chambers would have been in approximately stage 4 / 5 two days earlier at the time of temperature shift (Gandara and Drummond-Barbosa 2022) (Fig. 2A). Normally, follicle cells in a stage 4 / 5 egg chamber undergo an average of two mitotic divisions over one day as it matures to stage 6 / 7, after which follicle cells undergo a programmed developmental switch to endocycles up until stage 10A (Margolis and Spradling 1995; Calvi *et al*. 1998). Given that there were two extra endocycle genome doublings instead of the normal two mitotic divisions during the ∼24 hours from stage 4 / 5 to stage 6 / 7, these data also suggest that the duration of each induced endocycle is approximately equal to that of the mitotic cycles that they supplanted (Margolis and Spradling 1995). Moreover, the increased DNA content of these follicle cells at stage 10 suggests that starting endocycles early can override the undefined cell-autonomous mechanisms that arrest endocycles to limit follicle cell ploidy to 16C.

**Figure 2:**
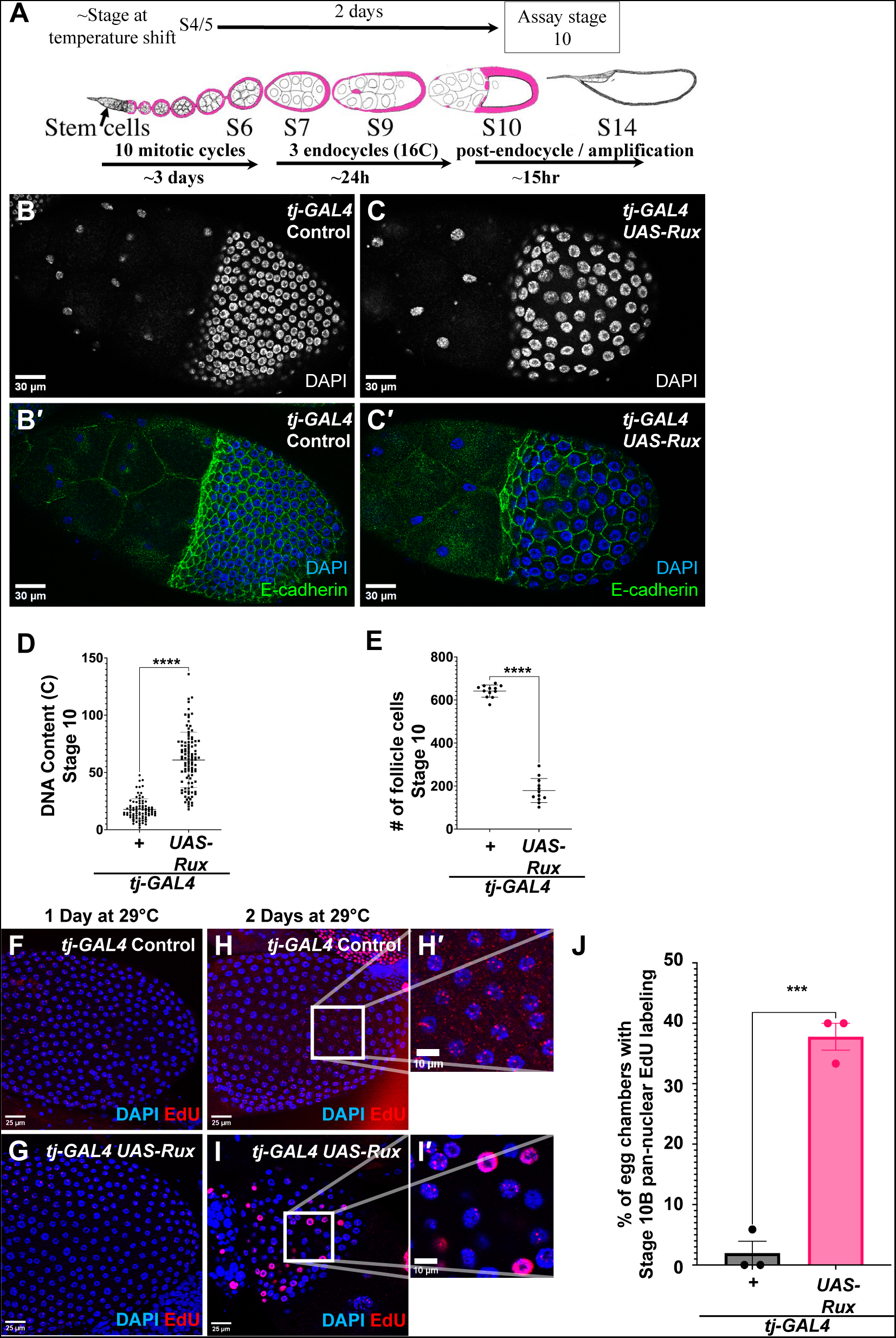
Premature endocycling increases follicle cell polyploidy and disrupts the endocycle-amplification transition. (A) An experimental timeline of the temperature shift experiments in relationship to the timing of egg chamber maturation. Select stages of oogenesis, follicle cell cycle transitions, and their approximate duration during oogenesis are indicated below. Follicle cells are pink. (B-C′) DAPI staining and immunolabeling against E-cadherin in control (B, B′) and Rux-expressing stage 10B egg chambers (C, C′); scale bar = 30 μm. (D) Quantification of DNA Content in control and Rux-expressing stage 10B egg chambers.; ****p < 0.0001; n = 85 nuclei in control and n = 104 nuclei in *UAS-Rux* across four biological replicates. (E) Quantification of number of follicle cells in control and Rux-expressing stage 10B egg chambers; ****p < 0.0001; n = 12 egg chambers in both across four biological replicates. (F-I′) DAPI staining and EdU labeling of DNA replication in control (F, H, H′) and Rux-expressing (G, I, I′) stage 10B egg chambers after one (F, G) or two (H-I′) days of incubation at 29°C; scale bar = 25 μm (F, G, H, I) or 10 μm (H′, I′). (J) Quantification of percent stage 10B egg chambers that had follicle cells with pan-nuclear EdU labeling in control and Rux-expressing ovaries after 2 days of Rux expression. ***p = 0.0003; n = 26 egg chambers for control and n = 19 egg chambers for *UAS-Rux* across three biological replicates.

We then examined the coordination of follicle cell cycles with later stages of oogenesis. in wild type, different follicle cells arrest endocycles at different times of stage 9-10A when they reach 16C. At the onset of stage 10B, follicle cells over the oocyte synchronously begin a specialized DNA replication program of developmental gene amplification (Fig. 2A) (Spradling and Mahowald 1980; Calvi *et al*. 1998). This selective gene amplification entails site-specific rereplication of specific loci that encode eggshell and other proteins, a process that is required for normal eggshell synthesis (Calvi and Spradling 1999; Calvi 2006; Hua and Orr-Weaver 2017). These rereplicating loci can be detected as distinct nuclear foci of EdU incorporation beginning in stage 10B (Calvi *et al*. 1998). EdU labeling of *tj-GAL4 GAL80^ts^ / +* control ovaries at 29°C showed that all follicle cells over the oocyte had the normal nuclear foci of EdU in stage 10B, indicating that the onset of amplification had the normal synchrony among follicle cells (Fig. 2F, H, H’) (Calvi *et al*. 1998; Calvi and Lilly 2004). Follicle cells that had been expressing Rux for 24 hours also had the normal transition to synchronous amplification in stage 10B (Fig. 2G). At the time of temperature shift one day earlier, most of these follicle cells would have already undergone the developmental switch to endocycles in stage 6 / 7, and therefore do not represent premature endocycling cells (Fig 2A). In contrast, two days after a temperature shift, the follicle cells in a stage 10B egg chamber are those that prematurely switched to endocycles in stage 4 / 5 (Fig. 2A). These premature endocycles resulted in some stage 10B egg chambers with a mixture of follicle cells that had either the normal amplification foci or pan-nuclear EdU incorporation indicative of continued genomic replication (Fig. 2I-J). The two-day delay in the onset of this phenotype suggests that starting endocycles early disrupts the later arrest of endocycle genomic replication and transition to developmental gene amplification.

### Premature endocycling inhibits border cell migration

During stage 9, most follicle cells migrate posteriorly to a position over the oocyte. Despite disruptions to cell structure and developmental amplification, this migration was normal in egg chambers with prematurely endocycling follicle cells, with most follicle cells achieving their normal position over the oocyte by stage 10A (Fig. 2C, C’). Unlike these main body follicle cells, a small group of follicle cells delaminates from the anterior epithelium and migrates posteriorly between the nurse cells (Montell *et al*. 1992; Spradling 1993; Montell *et al*. 2012; Mishra *et al*. 2019b). This collective migration event is initiated in early stage 9 when two specialized follicle cells, known as the polar cells, recruit 4-6 neighboring follicle cells to become border cells via JAK-STAT signaling (Montell *et al*. 1992; Silver and Montell 2001). After arriving at their final destination at the anterior of the growing oocyte in stage 10A, this migrating border cell cluster forms the sperm entry port known as the micropyle (King 1970; Bishop and King 1984; Horne-Badovinac 2020). Because these border cells have been a powerful model for collective cell migration during metastasis, we wished to investigate the effect of premature endocycling on them (Stuelten *et al*. 2018).

We identified border cells based on their elevated E-cadherin levels and measured their migration in both control and Rux-expressing stage 9-10A egg chambers two days after induction at 29°C (Oda *et al*. 1997; Niewiadomska *et al*. 1999). We normalized border cell migration to that of main body follicle cells on the surface of the same egg chamber during stage 9-10A. In controls, the migration of the border cell cluster between the nurse cells stayed on-pace with the anterior of the main body follicle cells, with both populations of follicle cells reaching the oocyte by the onset of stage 10A (Fig. 3A, A′, C). In contrast, Rux expression resulted in border cells lagging behind the main body follicle cells to varying extents (Fig. 3B-C). In some egg chambers, border cells had failed to even delaminate from the epithelium at the anterior pole of the egg chamber by stage 10 (Fig. 3C). Border cell migration was also impaired when premature endocycles were induced by *cyclin A* RNAi or *fzr* over-expression (Fig. S5A-C”). Moreover, using either *fru-GAL4* or *slbo-GAL4* drivers to induce high levels of Rux expression specifically in border cells after the developmental transition of follicle cells to endocycles in stage 6/7 did not impair border cell migration (Fig. S5D-I). These results indicate that the border cell migration defect is not because of unknown, off-target activities of the Rux protein, but rather a result of premature endocycling.

**Figure 3:**
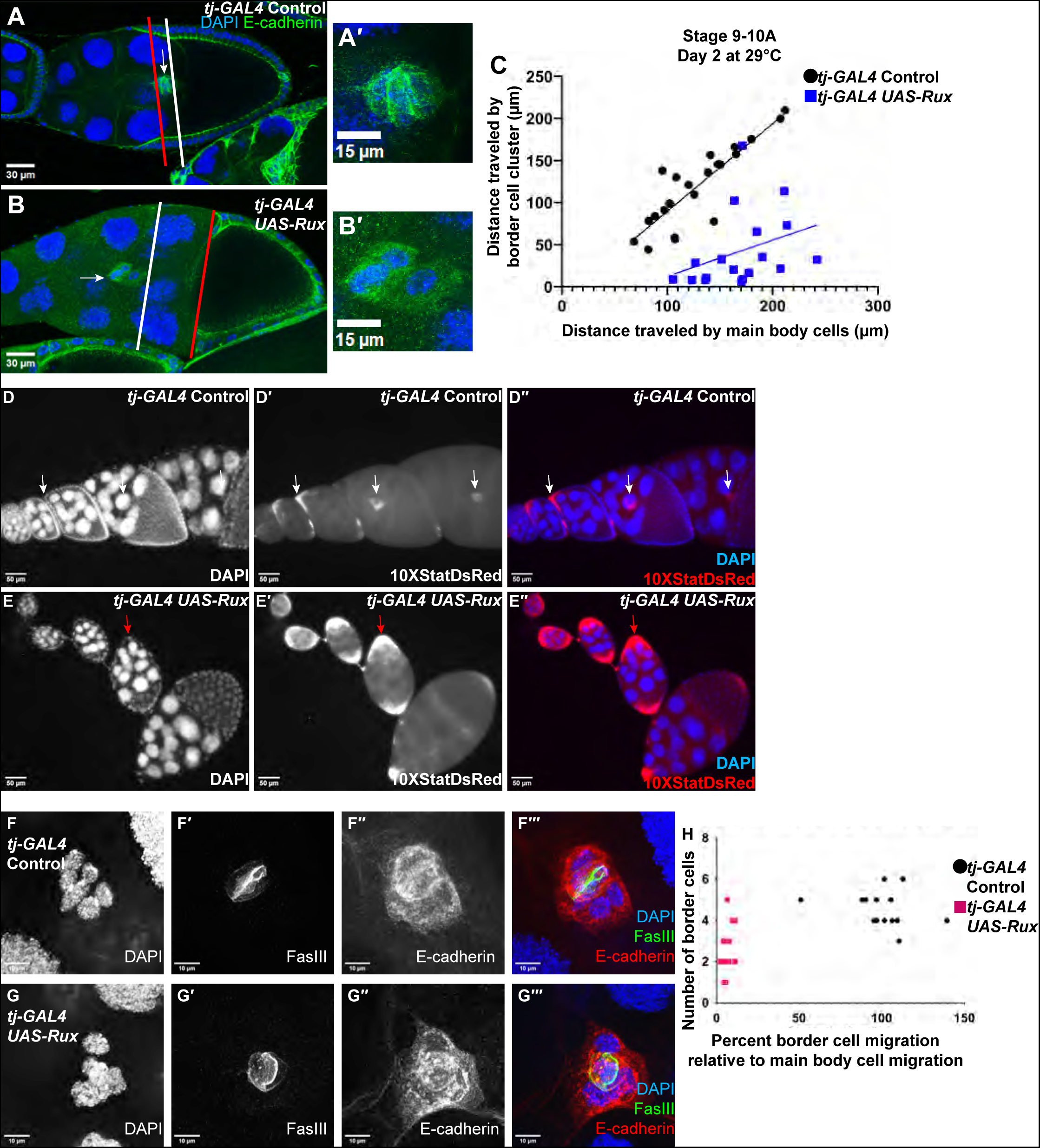
Premature endocycling inhibits border cell migration. (A-B′) DAPI staining and immunolabeling against E-cadherin in control (A, A′) and Rux-expressing (B, B′) egg chambers after two days of incubation at 29°C; scale bar = 30 μm (A, B) or 15 μm (A′, B′). White arrows indicate border cell clusters. White lines indicate extent of migration of border cell clusters, and red lines indicate extent of migration of main body cells. (C) Quantification of border cell migration at different times of stage 9 from control and Rux-expressing females after two days of incubation at 29°C across five biological replicates. Each dot represents one egg chamber with the distance migrated by its main body and border cells plotted on the x- and y-axis, respectively. (D-E′′) Expression of a JAK-STAT transcriptional reporter,*10XStatDsRed*, in control ovaries (D-D′′) and ovaries expressing *UAS-Rux* (E-E′′) after two days of incubation at 29°C. White arrows indicate anterior follicle cells and migrating border cell clusters in controls while red arrows indicate anterior follicle cells and delayed border cell clusters in Rux- expressing egg chambers. Scale bar = 50 μm. (F-G′′′) High magnification images of control (F-F′′′) or Rux-expressing (G-G′′′) border cell clusters after two days of incubation at 29°C. Clusters are visualized with DAPI staining and immunolabeling against FasIII and E-cadherin; scale bar = 10 μm. (H) Quantification of border cell number vs. border cell percent migration relative to main body cells in control and Rux-expressing ovaries. Each dot represents one egg chamber.

We used a *10XStat92E-DsRed* JAK-STAT transcriptional reporter to determine whether the failure to delaminate and migrate was because of a disruption of JAK-STAT signaling from polar cells to border cells (Silver *et al*. 2005; Bach *et al*. 2007). Both controls and *UAS-Rux* expressing ovaries had DsRed expression in follicle cells at the anterior and posterior ends of chambers, with the anterior signal narrowing to include only the border cell cluster by stages 9-10 of oogenesis, consistent with previous reports (Bach *et al*. 2007; Mallart *et al*. 2022) (Figure 3D-E”). The border cells in Rux-expressing ovaries had higher DsRed expression, which is likely due to higher copy number of the *10XStat92E-DsRed* reporter in these polyploid cells (Fig. 3E-E′′). These DsRed-positive border cell clusters had impaired migration, with some failing to delaminate from the epithelium, suggesting that these defects are not because of a failure in JAK-STAT signaling from polar cells to border cells.

To further investigate the border cell migration defect, we examined the morphology and number of cells in the border cell clusters. Labeling for the polar cell marker protein Fasciclin III (Fas III) showed that after two days of Rux expression most egg chambers had the normal two polar cells, including those in which border cells migrated slowly or failed to delaminate from the epithelium (Fig. 3F-G′′′) (Ruohola *et al*. 1991). Co-labeling against E-cadherin suggested that these migrating clusters had an overall normal structure and cell polarity (Fig. 3F-G′′′).

Rux-expressing egg chambers often had border cell clusters with a reduced number of border cells that failed to migrate normally (Fig. 3H). Given that it has been shown that a similar deficit of cells can inhibit border cell cluster delamination and migration, this observation suggests that the migration defect in these clusters is at least partially attributable to a reduced number of border cells (Silver and Montell 2001; Ghiglione *et al*. 2008). However, some clusters with the normal number of border cells also failed to migrate, suggesting that other properties of premature endocycling cells also inhibit migration (Fig. 3H).

### Premature endocycling impairs oocyte growth

We next examined whether premature follicle cell endocycles affected the growth of late-stage egg chambers. As mentioned previously, the overall morphology of stage 10B egg chambers from Rux-expressing females appeared similar to those from controls with main body follicle cells undergoing the normal migration posteriorly to over the growing oocyte (Fig 4A-D). One day after temperature shift stage 10B egg chambers from Rux-expressing females had oocytes of a similar size to controls (Fig. 4A, B, E). At the time of temperature shift one day earlier, the follicle cells in these stage 10B egg chambers would have been in stage 6/7 and already switched to developmental endocycles. In contrast, two days after temperature shift the oocytes in Rux-expressing stage 10B egg chambers was significantly smaller (Fig. 4C-E). The follicle cells in these stage 10B egg chambers represent those that had prematurely switched to endocycles in stage 4 / 5, suggesting that unscheduled endocycles impair oocyte growth. The egg chambers with smaller oocytes had a corresponding increase in the size of the nurse cell compartment (Fig. 4F). The ploidy of these nurse cells was not increased, suggesting that the enlarged nurse cell compartment was not because of enhanced nurse cell growth through additional endocycles (Fig. 4G). Labeling with the lipophilic dye Nile Red indicated that egg chambers with premature endocycling follicle cells had the normal accumulation of lipid droplets in nurse cells during stage 10 and their normal rapid transfer into the oocyte with the rest of the nurse cell contents during stages 11-12 (Fig. 4H-K). These results suggest that the two extra follicle cell endocycles had non-autonomous effects on oocyte growth before stage 11.

**Figure 4:**
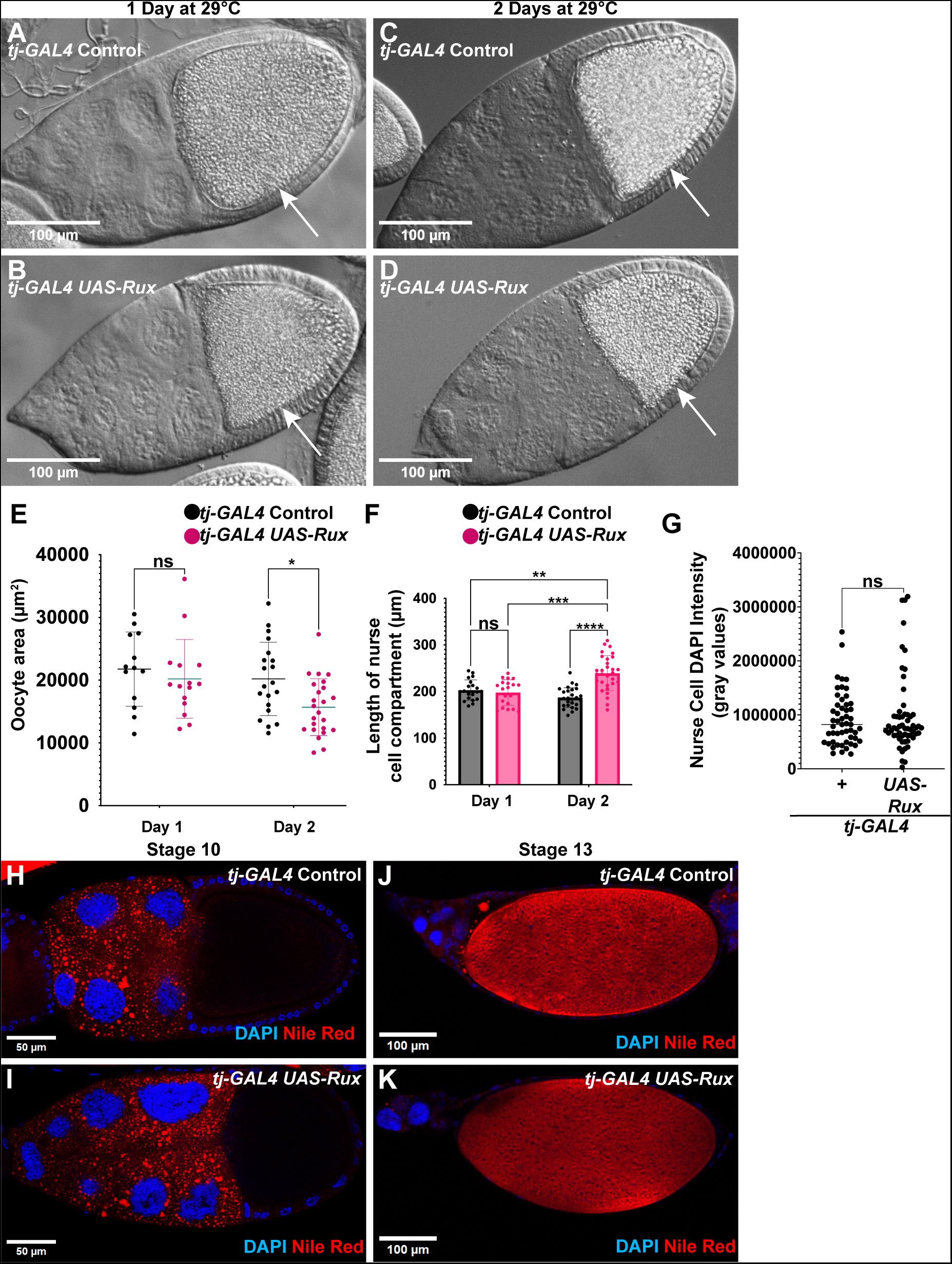
Premature endocycling of the follicular epithelium compromises germline growth. (A-D) Differential interference contrast images of control (A, C) and Rux-expressing (B, D) stage 10B egg chambers after one (A, B) or two (C, D) days of incubation at 29°C. The arrows indicate the posterior of egg chambers with the growing oocyte. scale bar = 100 μm. (E) Quantification of oocyte area in control and Rux-expressing stage 10 egg chambers.; *p < 0.05; for day one, n = 13 oocytes for control and n = 15 oocytes for *UAS-Rux* across three biological replicates; for day 2, n = 20 oocytes for control and n = 24 oocytes for *UAS-Rux* across five biological replicates. (F) Quantification of nurse cell compartment length; **p = 0.0021, ***p = 0.0002, ****p < 0.0001. For day 1, n = 20 egg chambers for control and UAS-*Rux* across three biological replicates; for day 2, n = 27 egg chambers for control and n = 30 egg chambers for *UAS-Rux* across five biological replicates. (G) Quantification of Stage 10 nurse cell ploidy. For control, n = 53 nurse cell nuclei; for Rux, n = 59 nurse cell nuclei. (H-K) DAPI staining of nuclei and Nile Red staining of lipid droplets in Stage 10B (H, I) or Stage 13 (J, K) egg chambers. Scale bar = 50 μm (H, I) or 100 μm (J, K).

### Premature endocycling disrupts eggshell synthesis and causes embryo lethality and female infertility

We evaluated whether the observed effects of premature endocycles on oogenesis would impact female fertility. It is known that defects in developmental gene amplification like those observed in Rux-expressing ovaries result in thin eggshells and a defective underlying vitelline membrane, ultimately causing dehydration and death of embryos (Calvi 2006). We therefore examined whether premature endocycles impaired eggshell synthesis. Some of the eggs laid by *UAS-Rux* females had thin and unhardened shells that resembled those from females with known defects in gene amplification, a phenotype that increased in frequency over one to three days (Fig. 5A, C, E; Fig. S6A, B) (Calvi 2006). Unlike wild type eggs, those from *UAS-Rux* females with abnormal shells were permeable to the dye Neutral Red, confirming a defect in eggshell and vitelline membrane synthesis (Figure 5B, D). Expression of Rux in females also resulted in reduced egg hatching, a phenotype that increased in severity over subsequent days of maternal Rux expression (Fig. 5F). This decrease in egg hatching correlated with the increase in collapsed, dehydrated eggs, suggesting that defects in vitelline membrane and eggshell synthesis were a significant cause of embryo lethality (Fig. 5E, F). Prolonged expression of Rux also had a negative effect on egg production during oogenesis. Finally, after three days of Rux expression there was a high frequency of ovarioles with degenerating stage 8-9 egg chambers, a phenotype diagnostic of an ovarian stress response known as the vitellogenic checkpoint (Fig. S6C, D) (Peterson *et al*. 2015; Serizier and McCall 2017). This timing suggests that in these ovaries a very early entry into endocycles during stage 1 / 2 triggers a general stress response three days later in stage 8. All together, these data indicate that premature follicle cell endocycles have deleterious effects on eggshell synthesis, embryo viability, and female fertility.

**Figure 5:**
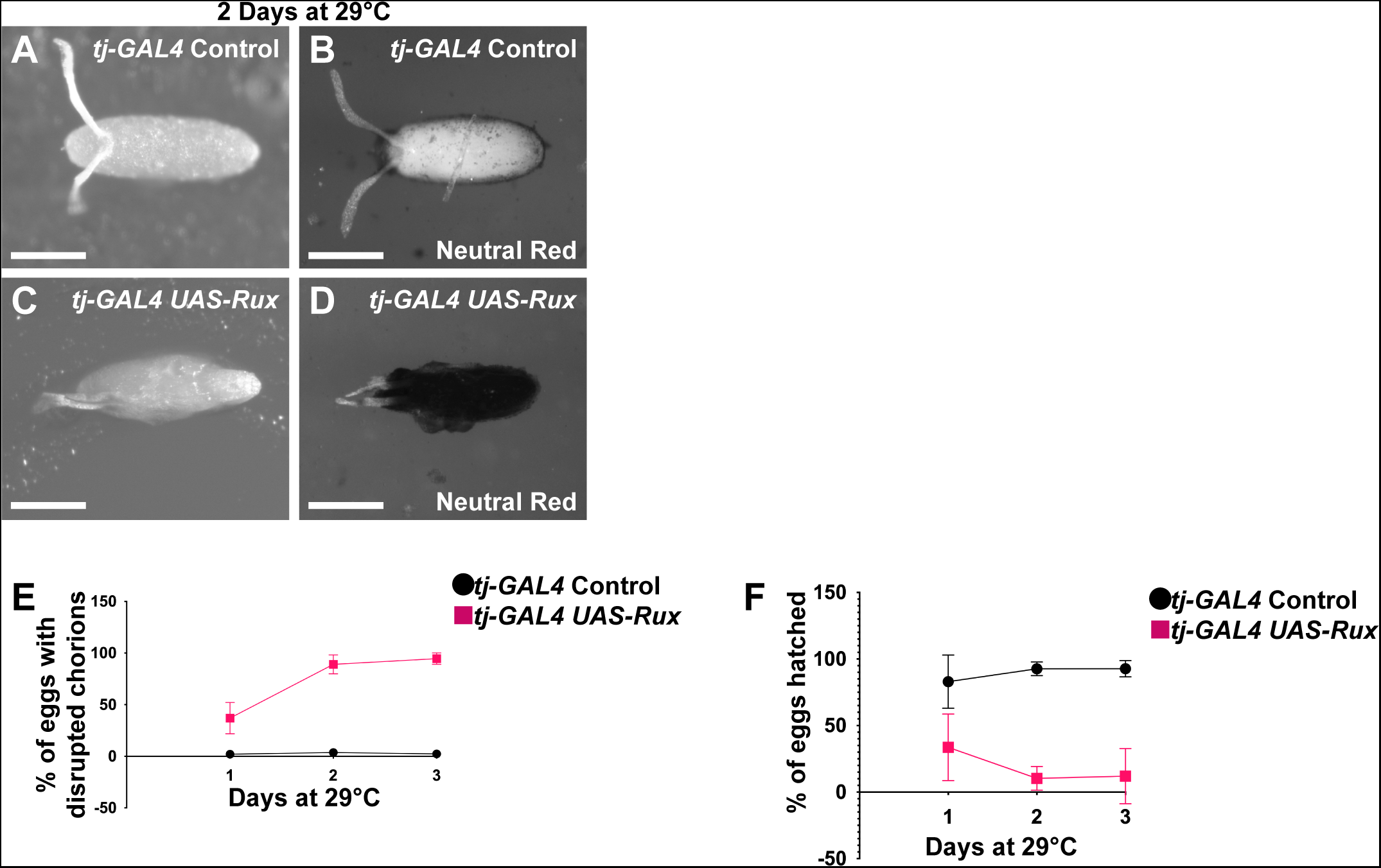
Premature endocycling disrupts eggshell synthesis and causes embryo lethality. (A-D) Brightfield images of eggs laid by control (A, B) and Rux-expressing females (C, D) after two days of incubation at 29°C; scale bar = 200 μm. Neutral red labeling is shown in grayscale (B, D). Quantification of percent of eggs with thin shells (E) and hatch rate (F) across three biological replicates.

## Discussion

There is a growing appreciation that cells can switch from mitotic cycles to endocycles in response to conditional signals, but the impact of this unscheduled switch on tissue growth and function is only beginning to be understood. We have found that premature follicle cell endocycles cause pleiotropic defects in oogenesis and severely compromise female fertility (Fig. 6). Although follicle cells normally switch to endocycles at stage 6 / 7 of oogenesis, inducing them to switch one day earlier in stage 4 / 5 resulted in aberrant cell and epithelial morphology, impaired oocyte growth, and perturbed the cessation of endocycles and transition to developmental gene amplification. These aberrant cell and tissue phenotypes are an inroad to defining the mechanisms by which cell growth and cell cycles are normally coordinated with stages of oogenesis. They also reveal the ways in which unscheduled endocycles can have deleterious consequences for tissue function that may apply across organisms, including in human disease.

**Figure 6:**
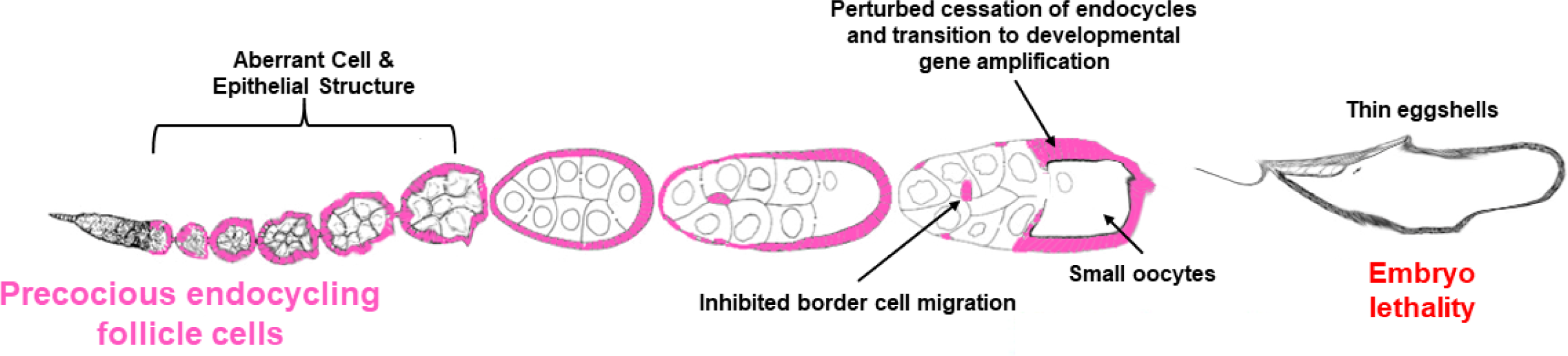
Premature endocycling causes pleiotropic defects in oogenesis. Premature entry of follicle cells (pink) into the endocycle causes aberrant cell and epithelial structure in early stages of oogenesis. These egg chambers with premature endocycles develop into later stages of oogenesis and have multiple defects including failure of border cell migration, disrupted endocycle to amplification transition, and non-autonomous effects on germline growth. The delay in these later defects after temperature shift indicate that they are the result a premature switch to endocycles in earlier.

Starting follicle cell endocycles early disrupted the later asynchronous, cell-autonomous arrest of endocycles during stage 9 and the synchronous onset of amplification at stage 10B (Calvi *et al*. 1998). Previous analyses of the endocycle to amplification transition has defined conserved mechanisms that regulate cell cycles and DNA replication during development. This transition is regulated by the E2F, RB, and Myb transcription factor families and the synchronous onset of amplification in stage 10B requires activity of Cyclin E / CDK2 (Calvi *et al*. 1998; Royzman *et al*. 1999; Bosco *et al*. 2001; Cayirlioglu *et al*. 2001; Beall *et al*. 2002; Beall *et al*. 2004; Calvi 2006). Further insight into the coordination of the endocycle to amplification transition with stages of oogenesis came from the finding that it requires downregulation of Notch signaling, which results in an obligate increase in the activity of the steroid hormone ecdysone and downstream expression of the transcription factor *tramtrack*, whose expression before stage 10 is further repressed by the micro-RNA *mir7* (Sun *et al*. 2008; Huang *et al*. 2013; Jia *et al*. 2015). Despite these key findings, much remains unknown about the molecular mechanism of endocycle arrest and the selective activation of amplification origins at precise times of oogenesis. Our finding that prematurely endocycling follicle cells can exceed their normal 16C DNA content by stage 10 suggests that their cell-autonomous endocycle arrest is not dependent on a strict genomic DNA counting mechanism. Some of these follicle cells increased their DNA content further through continued genomic replication past stage 10, which comprised the amplification of chorion and other genes that are required for eggshell synthesis. This disruption of amplification is likely the cause of the thin eggshells and embryo lethality that we observed, which phenocopies mutants that are known to impair gene amplification (Calvi 2006). A continued analysis of these induced endocycles may provide further insights into the molecular mechanisms that coordinate the timely transitions of cell cycle and DNA replication programs during development.

Although there was no evidence for a cell autonomous restriction on DNA content, there was a conservation of total DNA among all follicle cells. In wild type, the main body follicle cells in stage 10 collectively represented ∼10,256 haploid genome equivalents (641 cells * 16 C), whereas inducing premature endocycles resulted in egg chambers with fewer but more highly polyploid cells that represented a comparable ∼10,024 haploid genome equivalents (179 cells * 56 C). This DNA equivalency may simply be the result of swapping two mitotic cycles for two premature endocycles of equal duration. However, a similar conservation of genome equivalents has also been noted when damaged tissue is regenerated by unscheduled endocycles, a process termed compensatory cellular hypertrophy (CGH) (Tamori and Deng 2013; Cohen *et al*. 2018). Thus, there may be constraints on tissue growth that are proportional to genome equivalents, as previously suggested (Cohen *et al*. 2018).

Premature endocycles resulted in severely aberrant follicle cell morphology and an irregular follicular epithelium. These phenotypes contrast with the uniform cell and epithelial morphology of endocycling follicle cells from wild type ovaries. The basis for this difference between induced and developmental endocycles is currently unclear. One possibility is that induced and developmental endocycling cells differ in the pathways that coordinate their morphology with their hypertrophic growth to large cell size. For example, it is known that the activity of the Hippo-Yorkie growth pathway is coupled to multiple cell polarity and cell-cell junction protein complexes (Zheng and Pan 2019). Although premature endocycles resulted in aberrant cell shapes, sizes, and cell-cell contacts, we did not find clear evidence for a direct effect of induced endocycles on cell polarity. However, this last question was difficult to address given the highly aberrant cell morphology and a further investigation into the effects of unscheduled endocycles on cell polarity may be warranted.

The aberrant follicular epithelium may be related to the global effects of premature endocycles on egg chamber morphology. The deformations of the somatic follicle cell-germline interface before stage 6 suggests that premature endocycles may uncouple the coordinated growth of soma and underlying germline nurse cells, a process that is incompletely understood (Vachias *et al*. 2014; Giedt and Tootle 2023). Premature follicle cell endocycles also had non-autonomous effects on oocyte growth. Although stage 10 oocytes were smaller, the rapid transfer of lipid droplets and other nurse cell contents during stage 11 was normal, as was final egg size. These observations are consistent with a model wherein a somatic epithelium of fewer and more highly polyploid follicle cells impedes the slow transport of nurse cell contents into the growing oocyte before stage 11, but not the rapid transfer of nurse cell contents that occurs during stage 11 (Giedt and Tootle 2023). Smaller stage 10 oocytes may also be the result of a defect in vitellogenesis. This process entails paracellular transport of yolk protein across the follicular epithelium via a process that is enabled by a regulated increase in epithelial permeability beginning in stage 8, a state known as patency (Isasti-Sanchez *et al*. 2021; Riechmann 2021). Patency requires remodeling of cell-cell junctions through a mechanism that includes E-cadherin, whose labeling was altered in premature endocycling follicle cells (Horne-Badovinac and Bilder 2005; Isasti-Sanchez *et al*. 2021). Thus, the aberrant cell and epithelial morphology of induced endocycling cells may disturb the remodeling of cell-cell contacts that is required for patency and oocyte growth. More broadly, these results suggest that unscheduled endocycles may affect the coordinated growth and regulated permeability of epithelia in other tissues.

The highly penetrant negative effect of premature endocycles on border cell migration is likely due, in part, to a reduced number of border cells in the clusters, which has previously been shown to impair migration (Silver and Montell 2001; Silver *et al*. 2005; Ghiglione *et al*. 2008; Mishra *et al*. 2019a). Nonetheless, some clusters with the normal number of border cells also failed to migrate, indicating that there are other mechanisms that inhibit delamination and migration. Our observation of impaired collective border cell migration contrasts with evidence that polyploidy enhances migration of cancer cells *in vitro* and may promote collective cell migration during metastasis *in vivo* (Godinho *et al*. 2014; Wang *et al*. 2019; Mallin *et al*. 2023). An important implication of our observations, therefore, is that the effects of unscheduled endocycles on cell migration may be dependent on cell type and tissue context. It is also possible that oncogenic mutations in cancer cells may synergize with the polyploid state to enhance migratory properties.

Overall, our study has revealed that unscheduled endocycles can have severe pleiotropic effects on tissue growth and function. Further characterization of these effects will lead to a greater understanding of how endocycling, both scheduled and unscheduled, can promote tissue regeneration or contribute to disease.

## Materials and Methods

### Drosophila genetics

*w^-^; tj-GAL4 tub-GAL80^ts^ / CyO TwistGFP* flies were crossed to *w^1118^* (control) or *w^-^; UAS-Rux / SM6a* (experimental) flies, and their offspring were reared at 18°C during larval and early adult development. The resulting adult female offspring were fed wet yeast with males for 2 days at 18°C before being transferred to new wet yeast and shifted to 29°C for 1, 2, or 3 days (as noted in the text). The females and males were transferred to new wet yeast every 2 days during the experiments. This process was replicated in the *UAS*-*CycA^RNAi^*, *UAS-fzr*, and the *10XStatDsRed* experiments. Experiments with the *slbo-GAL4* and *fru-GAL4* border-cell-specific drivers were carried out at 25°C. The *w^-^; tj-GAL4 tub-GAL80^ts^* / *CyO TwistGFP* strain was a generous gift from L. Weaver. The *w^-^; UAS-Rux / SM6a* strain was obtained from the Bloomington Drosophila Stock Center (BDSC #9166). The *w^-^; 10XStatDsRed* strain was a generous gift from E. Bach. The *w^-^; slbo-GAL4 10XUAS-CD8GFP* strain was obtained from the BDSC (BDSC #76363). The *w^1118^; fru*-*GAL4* strain was obtained from the BDSC (BDSC #30027). The *w^-^; UAS*-*CycA^RNAi^* strain was obtained from the Vienna *Drosophila* Resource Center (VDRC v32421). The *w^-^; hsp70-GAL4 UAS-fzr / CyO; UAS-fzr* (*2XUAS-fzr*) strain is available upon request, and its components are available from the BDSC (BDSC #91688 for *UAS-fzr* on chromosome II and BDSC #91689 for *UAS-fzr* on chromosome III). All fly strains are also enumerated in the Reagent Table.

### Immunofluorescence microscopy

Ovaries were fixed and labeled as previously described (Schwed *et al*. 2002). For E-cadherin labeling, fixation buffers were supplemented with 1 mM CaCl_2_. The following antibodies and dilutions were used: rat anti-Shg (fly E-cadherin), 1:20 (DSHB DCAD2); mouse anti-Arm (fly β-catenin), 1:100 (DSHB N2 7A1); rabbit anti-phosphorylated histone H3 (pH3), 1:1000 (Millipore 06-570); mouse anti-Dlg1, 1:200 (DSHB 4F3); mouse anti-FasIII, 1:50 (DSHB 7G10); rabbit anti-GFP, 1:1000 (Invitrogen A-11122). EdU incorporation *in vitro* was for 1 hour as previously described (Calvi and Lilly, 2004). For lipid droplet detection, 10mg/mL Nile Red stock was diluted 1:500 in blocking buffer, and ovaries were incubated for 1 hour at room temperature with nutating, as previously described (Giedt *et al*. 2023). Asynchronous chorion gene amplification was defined as stage 10B egg chambers with both punctate and pan-nuclear EdU labeling of follicle cell nuclei. Egg chambers with more than one pan-nuclear EdU labeled nucleus were scored as “asynchronous.” For the dye exclusion assay, eggs were incubated in 1mL of 5 mg/mL of Neutral Red for 10 minutes, as previously described (LeMosy and Hashimoto 2000). Fluorescence micrographs were taken on a Leica SP8 confocal or a Leica DM5500 B. DIC images of stage 10B egg chambers were taken using a Leica DM5500 B. Brightfield images of laid eggs were taken using a Zeiss SteREO Discovery.V12. Nuclear DNA content was quantified using ImageJ/FIJI by measuring mean DAPI fluorescence per pixel in a nucleus, subtracting local mean background DAPI fluorescence per pixel, and then multiplying the difference by the number of pixels per nucleus to obtain the corrected total DAPI fluorescence intensity. To quantify the number of follicle cells per egg chamber, z-stacks of entire egg chambers were imaged, then DAPI-labeled nuclei were manually counted for the top part of the egg chamber where nuclei were most distinguishable from each other. This follicle cell count was then doubled to estimate the total number of follicle cells per egg chamber. The number of pH-positive follicle cells in early-stage egg chambers, total number of follicle cells in late-stage egg chambers, oocyte area, nurse cell compartment length, follicle cell diameter (along their longest axis), and border cell migration were quantified using ImageJ/FIJI.

### Eggshell scoring and egg hatch rate

Eggs from well-fed and mated females were collected on grape plates with a thin layer of yeast paste for 2-4 hours for each respective day of 29°C transgene induction. Eggs were scored as having “thin” chorions if they met the following criteria: 1) The eggshell appeared mucosal with no obvious follicle cell footprint pattern, 2) they had flaccid dorsal appendages, and / or 3) they were dehydrated and burst easily after a light touch with a paintbrush. The validity of these criteria and permeability of eggshell and vitelline membrane were independently confirmed by the Neutral Red staining described above. For hatch rates, eggs were lined up and incubated at 18°C to ensure no GAL4-driven transgene expression affected their viability. They were then scored for hatching over four subsequent days.

### Statistical Analysis

Data graphing and statistical analyses were carried out in GraphPad Prism. Data were tested for normality and homoscedasticity to determine correct analyses. Sample size (n) is defined as the total of independent observations / measurements used for the chosen analysis across multiple biological replicates. All averages shown are mean ± s.d., except for Figure 2J, which is mean ± s.e.m, and Figure 4G, which is median. Two-tailed Mann-Whitney tests were used to test for significance in Figs. 1F, S1E-F, 2D, 4E, and 4G. Figure 4E was corrected for multiple comparisons. A two-tailed Welch’s t-test was used to test for significance in Figs. 1G & 2E. A two-tailed unpaired t-test was used to test for significance in Fig. 2J. A Mixed Effects Two-Way ANOVA was used to test for significance in Fig. 4F.

## Data availability

Fly strains used are enumerated in the article. All fly strains used in the study are publicly available at the Bloomington Drosophila Stock Center, or upon request. All reagents used in this study are publicly available.

## Supporting information

Reagent Table

## Acknowledgements

We thank S. Brown and Y.T. Huang for experimental assistance. We are appreciative of the thoughtful feedback from S. Brown, L. Hesting, Y.T. Huang, P. Rangarajan, and C. Walczak, as well as from the editor and anonymous reviewers. We thank A. Kun and J. Powers of the IU Light Microscopy Imaging Center. Special thanks to D. Montell for advice and E. Bach and L. Weaver for fly strains

## Author Contributions

H.C.H. performed experiments and statistical analyses. Both authors wrote and edited the manuscript. B.R.C. secured funding and provided mentorship and supervision.

