## Supplemental Figure Legends for "Premature endocycling of *Drosophila* follicle cells causes pleiotropic defects in oogenesis"

### Supplemental Data Figure Legends

**Figure S1: Rux expression induces premature endocycling.** DAPI staining, EdU labeling of DNA replication, and immunolabeling against pH3 in control (A, A′, C, C′) and Rux-expressing (B, B′, D, D′) ovarioles after one (A-B′) or two (C-D′) days of incubation at 29°C. Yellow arrows indicate pH3-labeled follicle cells and pink arrows indicate EdU-labeled follicle cells. Stages of oogenesis (S) and germarium (G) are indicated. Scale bar = 30 µm. (E-F) Quantification of the number of phospho-Histone H3 (pH3) labeled cells in control and Rux-expressing egg chambers after one (E) and two (F) days of incubation at 29°C. ****$p<0.0001$.

**Figure S2: Premature endocycling results in aberrant Armadillo labeling.** (A-B″) DAPI staining and immunolabeling against Armadillo in cross-section images of control (A-A″) or Rux-expressing (B-B″) egg chambers after two days of incubation at 29°C; scale bar = 20 µm. Stages (S) of oogenesis and germarium (G) are indicated.

**Figure S3: Effects of Rux expression on Dlg1 localization.** (A-B″) DAPI staining and immunolabeling against Dlg1 in control (A-A″) and Rux-expressing (B-B″) stage 4/5 egg chambers after two days of incubation at 29°C. White arrows indicate single follicle cells. Scale bar = 10 µm. (C-D″) DAPI staining and immunolabeling against Dlg1 in control (C-C″) and Rux-expressing (D-D″) stage 10B egg chambers after two days of incubation at 29°C; scale bar = 30 µm.

**Figure S4: *CycA* RNAi and *fzr* over-expression iECs have aberrant cell and epithelial morphology.** (A-C″) Stage 4-5 egg chambers in control (A-A″), *UAS-CycA*$^{RNAi}$ (B-B″) or *UAS-fzr* expressing (C-C″) ovaries two days after shifting to 29°C. All three genotypes had *tj-GAL4 tub-GAL80*$^{ts}$. Ovaries were stained with DAPI and immunolabeled against E-cadherin to visualize cellular and epithelial structure defects. Scale bar = 10 µm.

**Figure S5: Premature endocycles cause border cell migration defects.** (A-C″) Premature endocycles induced by *CycA* knockdown or *fzr* overexpression cause border cell migration defects. Stage 10A egg chambers in control (A-A″), *UAS-CycA*$^{RNAi}$ (B-B″) or *UAS-fzr* expressing (C-C″) ovaries after two days of expression with *tj-GAL4 tub-GAL80*$^{ts}$. Ovaries were stained with DAPI and immunolabeled against E-cadherin to visualize border cell migration. White arrows indicate border cell clusters. White lines indicate extent of migration of border cell clusters, and red lines indicate extent of migration of main body cells. (D-I)

Expression of *UAS-Rux* after the switch to developmental endocycles does not impair border cell migration. (D-D′) The expression domain of *slbo-GAL4* in the ovary was visualized with a *10XUAS-CD8GFP* transgene and immunolabeling against GFP. (E-F) Stage 10A egg chambers in *slbo-GAL4 10XUAS-CD8GFP* control (E) and *slbo-GAL4 10XUAS-CD8GFP / UAS-Rux* (F) ovaries were stained with DAPI and immunolabeled against E-cadherin and GFP to visualize border cell migration. (G-G′) The expression domain of *fru-GAL4* in the ovary detected with a *10XUAS-CD8GFP* transgene and immunolabeling against GFP. (H-I) An egg chamber from *fru-GAL4 10XUAS-CD8GFP* control (H) and *fru-GAL4 10XUAS-CD8GFP / UAS-Rux* (I) ovaries were stained with DAPI and immunolabeled against E-cadherin and GFP to visualize border cell migration. White arrows indicate border cell clusters. White lines indicate extent of migration of border cell clusters, and red lines indicate extent of migration of main body cells.

**Figure S6: Three days of Rux expression severely disrupts eggshell synthesis and activates the vitellogenic checkpoint.** (A, B) Brightfield images of eggs laid by control (A) or Rux-expressing (B) females after three days of incubation at 29°C. (C, D) DAPI staining in control (C) or Rux-expressing (D) ovarioles after three days of incubation at 29°C. Scale bar = 200 μm (A, B), or 100 μm (C, D).
