## Supplementary figures and images for "Premature endocycling of *Drosophila* follicle cells causes pleiotropic defects in oogenesis"

### Supplemental Figures

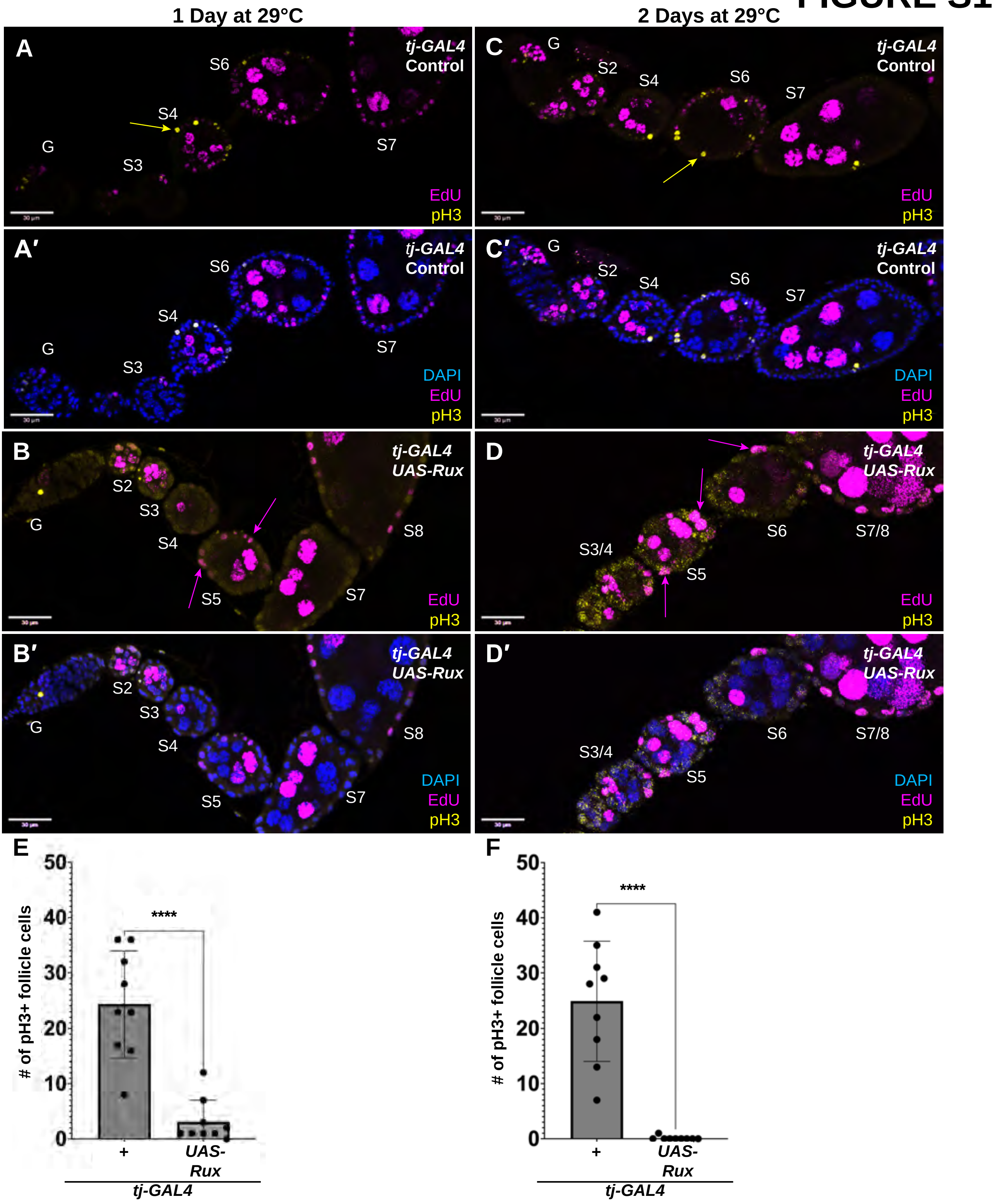

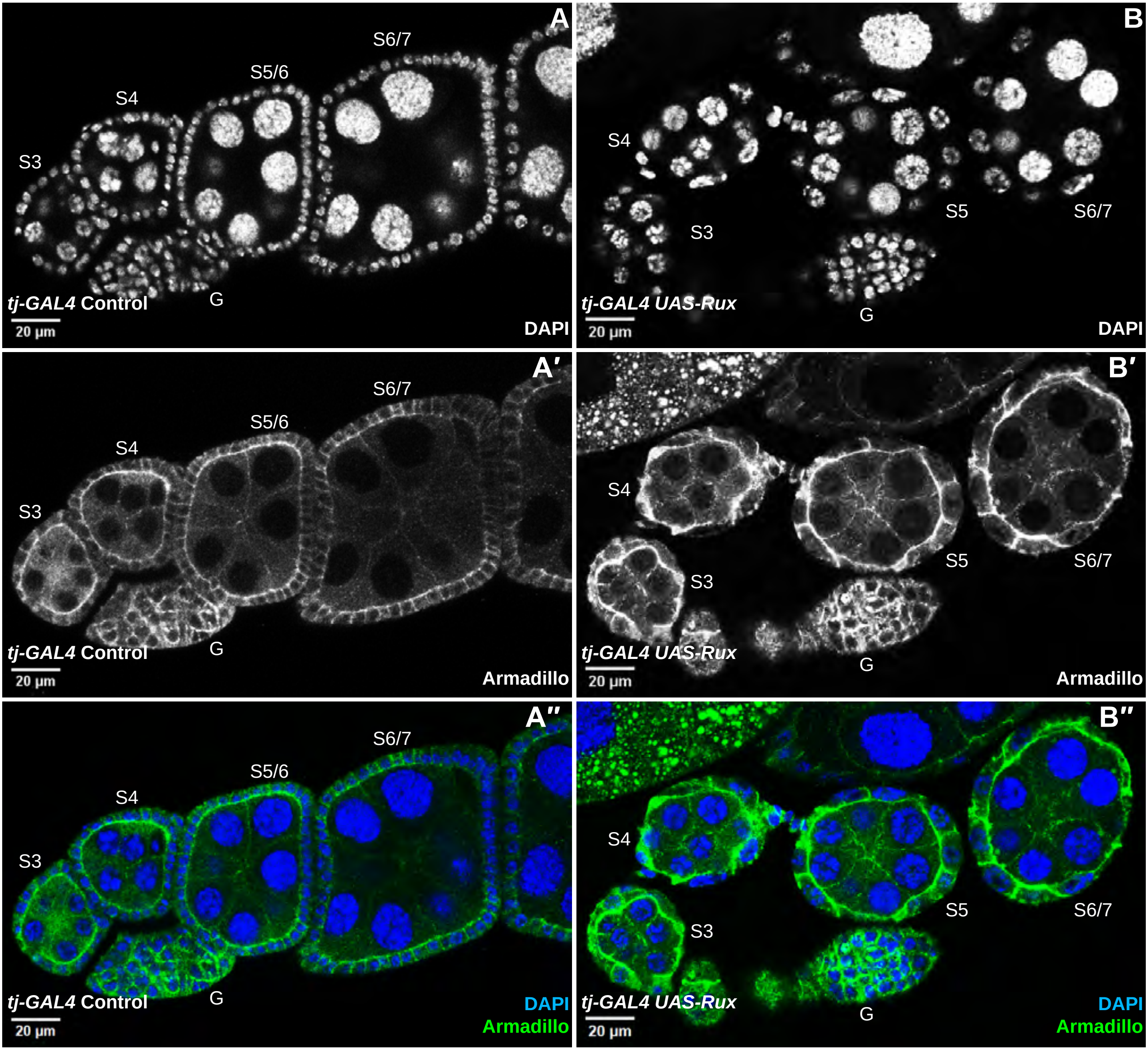

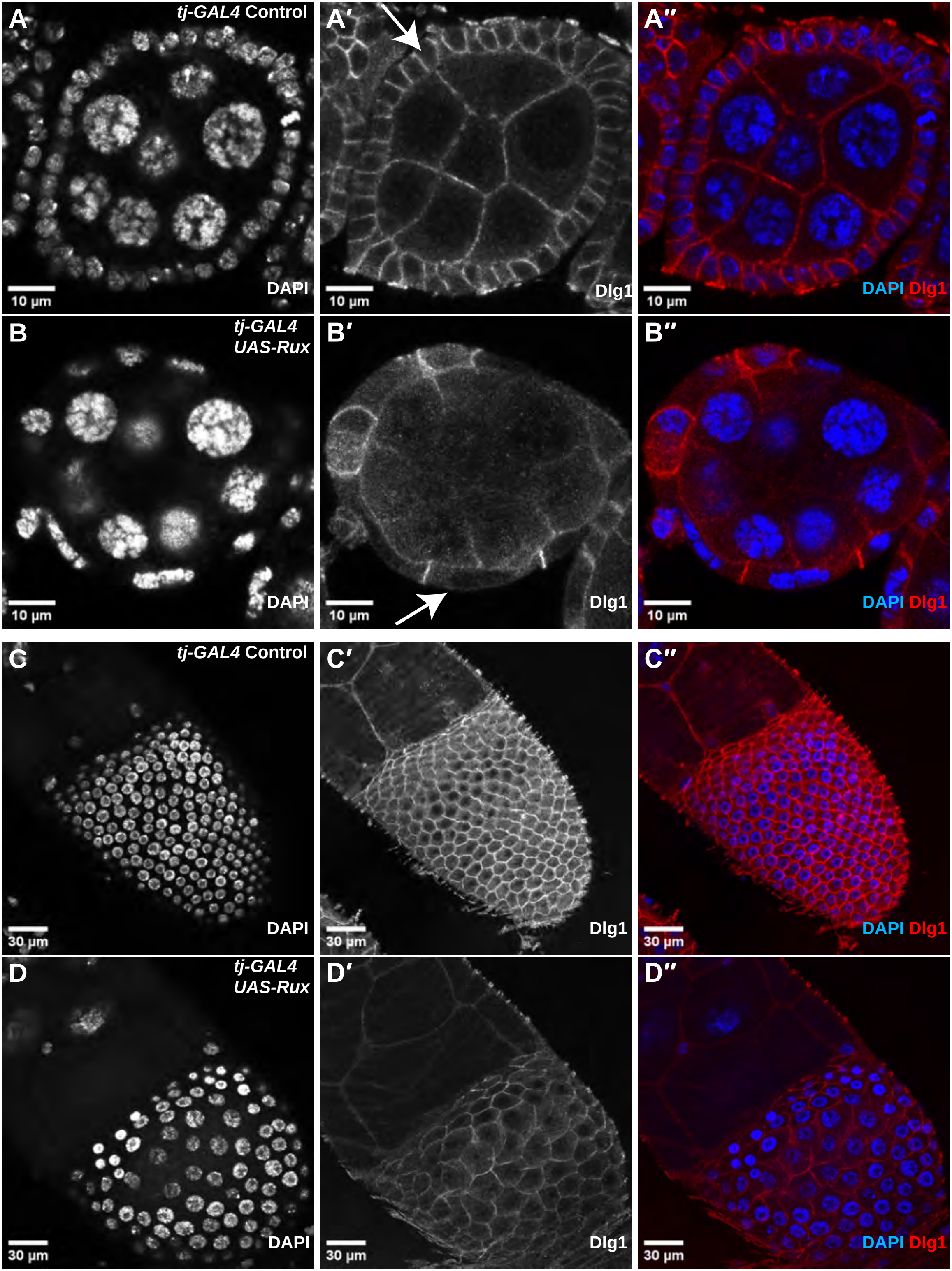

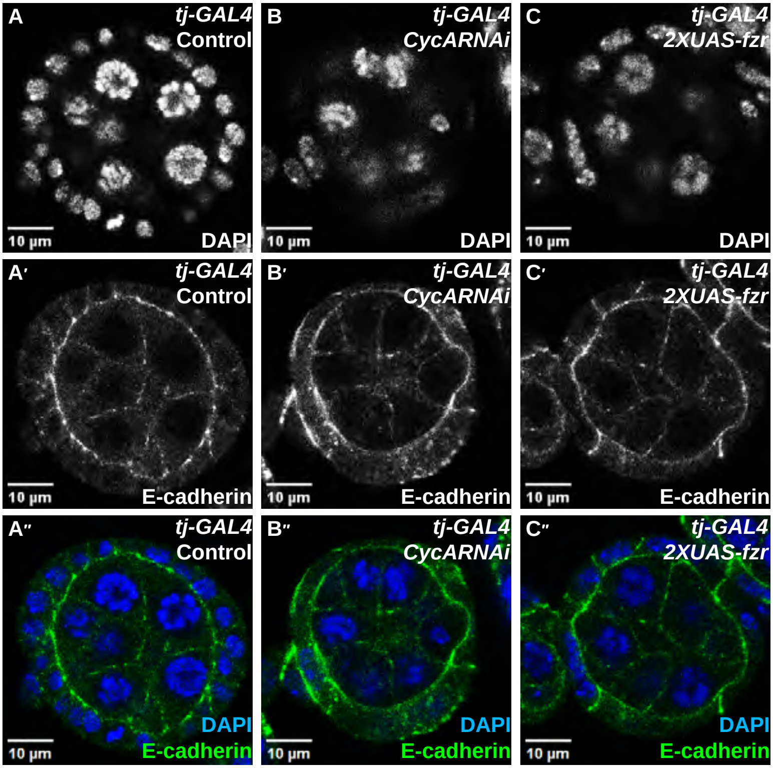

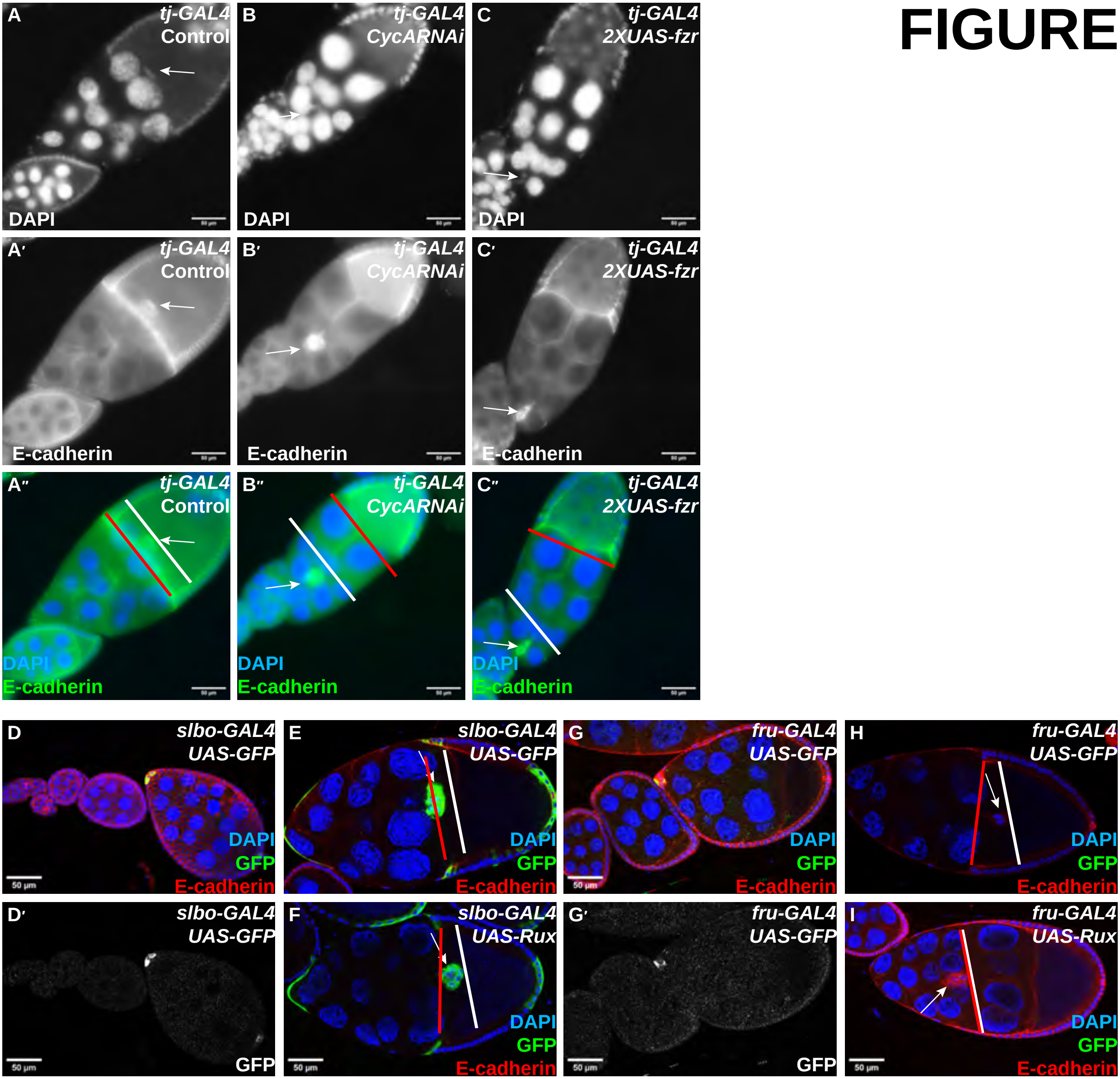

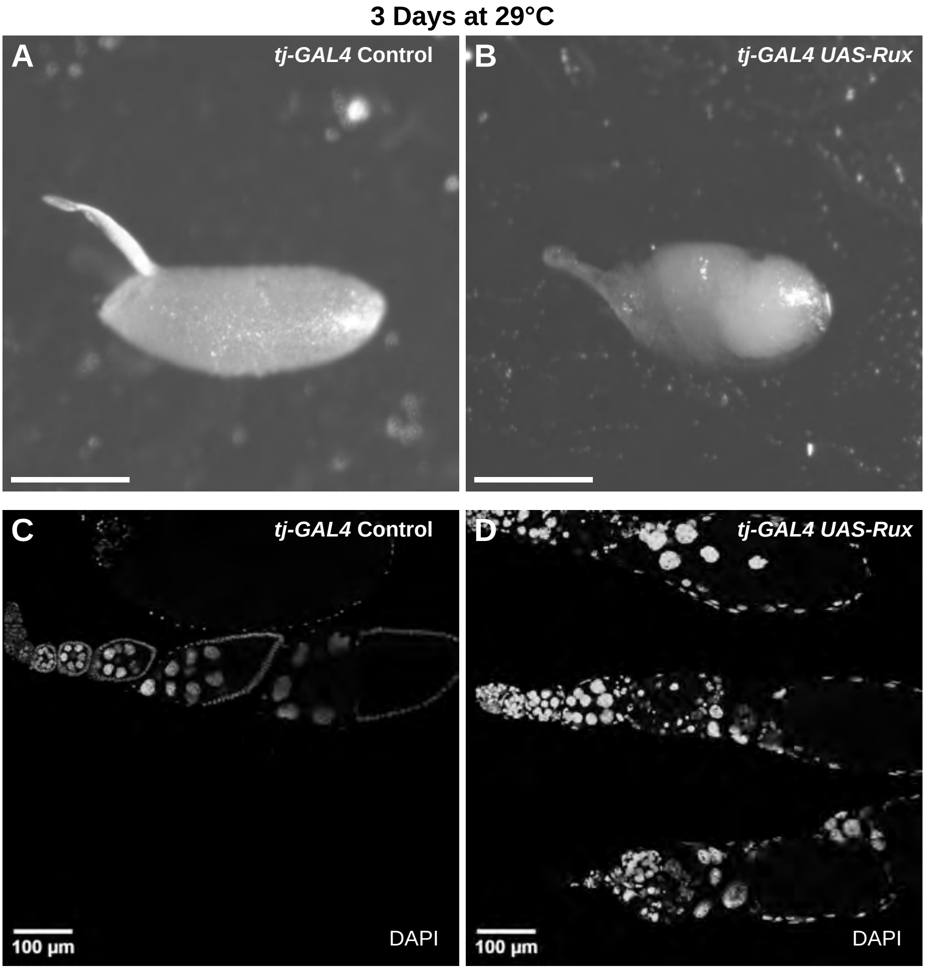
